# Variation in scion water relations mediated by two newly released Geneva series rootstocks

**DOI:** 10.1101/2021.07.20.453132

**Authors:** E. Casagrande-Biasuz, L.A. Kalcsits

## Abstract

Dwarfing rootstocks are used to control tree vigor allowing for increased densities that increase apple production. Although there is considerable variation among rootstocks in dwarfing capacity, the mechanisms by which rootstocks affect vigor in apple scions remains unclear. Here, ‘Honeycrisp’ apple growth and water relations were compared among three rootstocks; M-9 as the industry standard and two less studied Geneva series rootstocks; G.87 and G. 814 in Washington, USA. Trees were acquired from a commercial nursery and planted in 2017. In 2018 and 2019, scion physiological, isotopic and morphological traits were measured to better understand the link between rootstock-driven vigor and physiological traits. Rootstock affected scion shoot growth (P <0.001), stomatal conductance (P< 0.01) and stem water potential (P <0.001). Rootstocks with low vegetative vigor like M.9 also had lower stomatal conductance and enriched leaf δ^13^C and δ^18^O isotope composition. Plant growth was positively correlated with stomatal conductance and stem water potential. Rootstocks also affected plant water status and net gas exchange. Here, we report an association between rootstock-induced vigor and scion physiological traits such as gas exchange, stem water potential, and leaf carbon and oxygen isotope composition. This research has implications for the understanding of the mechanisms of dwarfing by rootstocks in apple.

## INTRODUCTION

High density apple orchard systems are made possible through the adoption of dwarfing rootstocks (Webster, 1995). Dwarfing rootstocks redirect photosynthate allocation away from shoots and branches and into fruit production (Atkinson and Else, 2001) producing higher quality fruit and higher yields. Although the scion produces the desirable cultivar we eat, the rootstock supports its growth (Gregory et al, 2013). Rootstocks can also be used to improve water and nutrient uptake (Tworkoski and Fazio, 2015), reduce disorders (Blanco et al.; unpublished) contributing to enhanced economic and environmental sustainability. Although dwarfing has been extensively characterized in different apple cultivars, the underlying physiological mechanisms contributing to this dwarfing effect remain unclear (Simon and Chu, 1984; Marguerit et al, 2012; Foster et al, 2017; Xu and Ediger, 2021). Some theories attribute the dwarfing mechanism of rootstocks to changes to the supply of water from the rootstock to the scion (Atkinson, 2003). A decline in hydraulic conductance may induce important effects on the scion which affect shoot growth (Atkinson et al., 2003). More conservative plants have lower water potential that is influenced by limitations to water movement from the soil and within the plant which produces a strong decrease in stomatal conductance and consequently, leads to lower photosynthetic rate and growth of trees (Jones, 1998; Brodribb et al., 2003).

Here we tested the association between dwarfing in three commercially available rootstocks, water relations, and downstream effects on net gas exchange and consequently, C and O isotope composition.

## MATERIAL AND METHODS

### Site description and experimental design

The experiment was conducted using ‘Honeycrisp’ grafted on three rootstocks at Washington State University Sunrise Research Orchard in Rock Island, WA (47°18’35.6”N 120°03’59.5”W). The soil is a shallow sandy loam soil. Rootstocks M.9-T337 (M.9), G.814 and G.87 were used for this study. The trees were planted in 2017 and arranged in a completely randomized design (N = 3) with eight trees per replication.

### Physiological and morphological measurements

Stomatal conductance, transpiration rate and net CO2 assimilation were measured on two sun-exposed fully mature leaves per replicate between 10am and 12pm using LI-6400XT infrared gas analyzer (Li-COR, Lincoln, NE, USA). Reference CO_2_ concentration was set to 400 ppm, leaf temperature at 25 °C, and photosynthetic photon flux density inside the chamber was set to 1,500 μmol m^−2^ s^−1^. Midday stem water potential was measured on two leaves per replicate using a Scholander System Pressure Chamber Instrument (PMS Instrument Co., Albany, OR, USA) at solar noon. Fully mature expanded leaves were selected from inside the canopy closest to the base of the tree and allowed to equilibrate with the stem in a silver reflective bag for at least 90 minutes. These measurements were made monthly for four months and values were averaged to produce an average summer value for each replicate. The measurements for plant growth took place in concomitance with physiological measurements from June to August. Shoot growth was measured as shoot length (cm) at the end of the growing season for three shoots on two trees in each replication.

### Isotope analysis

For isotope biomass analysis, six shoots with leaves were collected from the upper-middle portion of the tree from both sides from three trees per replication. The stems were sampled on three different times from June to August. The samples were collected in the morning, leaves and shoots were detached and separated in different paper bags for each replicate. All samples were then brought back to the lab and placed in a chamber with constant air flow at 25 °C until reached a stable weight. Once dry, shoots and leaves were used for carbon and oxygen isotope analysis.

#### 1. Carbon isotope composition (δ^13^C)

Dry leaves and stems were well-mixed and ground in fine powder using a VWR Homogenizer (VWR, Radnor, PA). Fine powder from leaves and shoots were weighed with the same precision analytical balance and placed into 5mm x 9mm tin capsules (Costech Analystycal Technologies, Inc., Valencia, CA, USA). The capsules were shipped for analyses at the Stable Isotope Core Laboratory at WSU and then analysed for δ^13^C using a Delta XP ThermoFisher isotope ratio mass spectrometer coupled with an elemental analyzer. Carbon isotope ratios were determined using the standard Pee Dee Belemnite (PDB) and the values reported in “delta” notations as *δ* values in parts per thousand (‰) (Farquhar et al., 1989a) as described in Eq. 1 and Eq. 2:

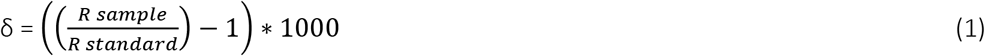

*Where R sample is the ratio of the heavy over the lighter isotope composition of the sample divided by the heavy over the lighter isotope composition of the standard (R standard)*.

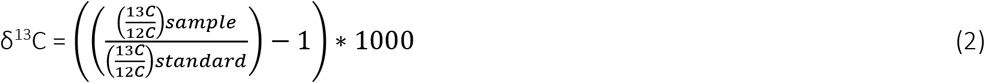

Following Eq. 1, to determine the carbon isotope composition of plant material, we calculate the ratio of the heavy (^13^C) over the lighter (^12^C) isotope value of the sample and divided by the ratio of the heavy (^13^C) over the lighter (^12^C) isotope value of the standard (Eq. 2). When comparing with the standard values or among other samples, more positive values of δ implies in higher concentration of heavy isotopes (enriched) whereas more negative values indicate higher concentration of lighter isotopes (depleted).

#### 2. Oxygen isotope composition of leaf biomass (δ^18^O_biomass_)

The same samples that were used for carbon isotope analysis were used for oxygen isotope analysis of leaf biomass. The samples then were weighted between 0.6-0.8mg of ground leaf tissue was weighed into 4×6mm silver capsules using a precision analytical balance (XSE105 DualRange, Mettler Toledo, Greifensee, Switzerland). Leaf tissue was analyzed using a continuous-flow pyrolysis using TC/EA interfaced with an IRMS (Delta Plus, ThermoFinnigan, Bremen, Germany) through a continuous flow device (Conflo-III, ThermoFinnigan,Bremen, Germany) at the Stable Isotope Core Laboratory at Washington State University, United States. Oxygen isotope ratios of each sample were then determined, and the values reported in “delta” notations as δ values in parts per thousand (‰) (Barbour, 2007) as described in Eq. 3 and Eq. 4:

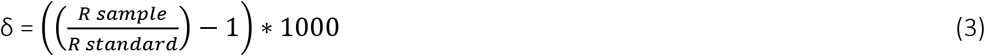

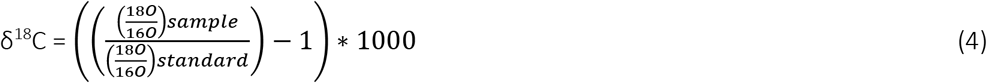

Where ^18^O/^16^O_sample_ is the enriched oxygen over the light oxygen in the samples and ^18^O/^16^O_standard_ is the known isotopic composition from the Vienna-Standard Mean Ocean Water (VSMOW) international standard commonly used for oxygen isotope analysis which present an isotope ratio of 2.0052 x 10-3 (Gonfiantini, 1984).

#### 3. Oxygen isotope composition of leaf water (δ^18^O_water_)

After mid-day water potential and leaf gas exchange measurements, following the sampling method described for the biomass analysis, leaves and stems were then sampled and inserted in plastic insulated vials (DWK Life Sciences Wheaton™ Liquid Scintillation Vials, Thermo Fisher Scientific, Waltham, MA) sealed with parafilm and placed immediately on ice to be transported and stored until water extraction. Water from samples was extracted using the distillation process described by Vendramini & Sternberg (2006). The δ^18^O from the samples water then, were determined by on-line pyrolysis method equilibration technique. The isotopic analysis of oxygen in plant and water use carbon monoxide as the target gas after pyrolyzing the sample by a modified procedure. The oxygen isotope composition in the water extracted from the samples will be calculated and reported as δ values in parts per thousand (‰) and then illustrated as the enrichment above source water (Flanagan and Ehleringer, 1991, Barbour et al, 2004; Flanagan & Farquhar, 2014).

### Statistical Analysis

Data were analyzed using SAS (SAS Institute, Cary, NC) via an analysis of variance (ANOVA) with rootstock as the main factor for shoot growth, stomatal conductance, transpiration rate, midday stem water potential. Mean separation tests were performed using Fisher’s LSD test with 95% confidence of significance. Figures were prepared using Origin2021 Data Analysis and Graphing Software (OriginLab Corporation, MA, USA).

## RESULTS AND DISCUSSION

Both G.814 and G.87 had greater shoot length than M.9. Similarly, M.9 had a more negative seasonally-averaged stem water potential when no water limitations were present. The midday stem water potential measured in this well irrigated orchard ranged from −0.87 to −0.98 MPa (Table 1). These observations follow similar patterns observed for different rootstocks in apple (Sarita et al., 2013; Xu et al., 2021) has been previously reported for apple rootstocks. Like stem length, stem water potential was the lowest for M.9. Stem water potential for G.814 and G.87 were not different. Midday stem water potential has been reported to be lower in more dwarfing rootstocks in other studies (Naor et al., 1995). Olien and Lakso (1986) observed a high correlation between stem water potential and rootstock than stem water potential and overall tree size suggesting that rootstocks effects on midday stem water potential may be independent of tree vigor. However, in this study, the two rootstocks with the most vigor also had the highest stem water potential.

**Table 1.** Shoot length (cm), stem water potential (Mpa), photosynthetic rate (CO_2_ m^−2^s^−1^), stomatal conductance (mmol H_2_O m^−2^s^−1^), transpiration rate (mmol H_2_O m^−2^ s^−1^) and carbon isotope composition in leaves (‰) measured on ‘Honeycrisp’ grafted onto M.9, G.814 and G.87 rootstock cultivars. Different letters denote significant differences among columns determined using a Fisher mean of separation test (alpha = 0.05). ** and * indicate significance in differences among means at p-values <0.01 and <0.05, respectively.

| *Measurements* | **Rootstock** |  |  |  |
| --- | --- | --- | --- | --- |
|  | **M.9** | **G.814** | **G.87** |  |
| **Shoot length (cm)** | 27.6 a | 49.2 b | 42.2 b | ** |
| **Stem water potential (MPa)** | -0.98 b | -0.91 a | -0.87 a | * |
| **Net $\text{CO}_2$ assimilation** | 15.1 a | 16.4 ab | 17.1 b | * |
| **Stomatal conductance** | 0.18 a | 0.19 ab | 0.21 b | * |
| **Transpiration rate** | 4.2 | 4.5 b | 5.0 b | * |
| **Leaf $\delta^{13}\text{C}_{\text{biomass}}$ (‰)** | -25.6 a | -25.8 ab | -26.1 b | * |
| **Leaf $\delta^{18}\text{O}_{\text{biomass}}$ (‰)** | 23.91 b | 24.44 ab | 24.96 a | * |
| **Leaf $\delta^{18}\text{O}_{\text{water}}$ (‰)** | 8.93 a | 6.78 a | 8.31 a | ns |

Since the rate of water flow through the plant is a function of the difference in soil water potential and the sum of plant resistance during uptake from soil and distribution to leaves, changes in water potential can be linked to changes in transpiration gradients in leaves (Shackel et al., 1997). Stomatal conductance is a strong regulator of overall plant transpiration (Li et al., 2002). Rootstocks significantly differed in stomatal conductance under the same water availability. M.9 had significantly lower stomatal conductance and, consequently, CO2 assimilation, compared to the more vigorous rootstock G.87. These findings are consistent with those reported previously (Stutte et al., 1994; Naor et al., 1995; Sarita Devi, 2013; Valverdi and Kalcsits, 2021). G.87, the most vigorous rootstock, had significantly higher transpiration rates compared to more dwarfing rootstock M.9.

Carbon isotope composition was affected by rootstock genotype (Table 1). Carbon isotope composition ranged from −26.1 to −25.6 ‰. Leaf carbon isotope composition of M.9, the least vigorous rootstock, was higher than the most vigorous rootstock, G.87. G.814 was not statistically different than M.9 and G.87. Oxygen isotope composition of leaf biomass was also significantly different among rootstocks. Opposite of δ^13^C, δ^18^O was the lowest for M.9 and the highest for G.87. However, δ^18^O for water extracted from leaves was not affected by rootstock. Stomatal conductance has been closely linked to carbon (Kalcsits et al. 2021; Johnson et al., 1990; Valverdi and Kalcsits, 2021) and oxygen isotope composition of leaves (Farquhar et al., 1989b; Barbour et al., 2000).

Carbon isotope composition can, therefore, be impacted by environmental water limitations and differences in hydraulic traits that affect stomatal conductance and gas exchange between the inner leaf and the atmosphere. In general, water conservative plants present higher carbon isotope values (Liu et al, 2012).

The effect of rootstock on water relations can be observed at physiological level that affects gas exchange including both water loss and CO2 assimilation. These affects are imprinted in the accumulation plant carbon biomass indicated by vigor as well as in the isotope composition in that newly assimilated carbon. In non-fruiting apple trees where stem and leaf biomass represents most of the carbon gain by the tree, there was a strong correlation between δ^13^C of leaf biomass and gas exchange traits (Figure 1). Similar, albeit less robust, relationships were observed between δ^18^O and gas exchange traits. Stem water potential was also negatively correlated with leaf δ^13^C. Therefore, higher carbon isotope composition of rootstock M.9 indicates inherent water limitations that are transferred from the rootstock to the scion. Since stomatal regulation is responsible for controlling stem xylem water potential, higher water limitations may be a consequence of reduced hydraulic conductance in dwarfing rootstocks. Hydraulic resistance in rootstocks may reduce the capacity of photosynthetic rate at scion level which may then translate to reduced growth (Else et al., 2000; Atkinson et al., 2003). In this study, rootstock also had a significant effect on tree growth ranging from 20 cm in the least vigorous M.9 to almost 50 cm for G.87 and G.814. Rootstock M.9 had significantly smaller shoot growth compared to G.87 and G.814. We observed that the one of the main influences of dwarfing rootstock is the capacity to produce lower biomass compared to vigorous rootstocks. It is evident the relationship of hydraulic resistance to lower water status and reduced shoot growth in M.9 compared to more vigorous rootstocks.

**Figure 1.**
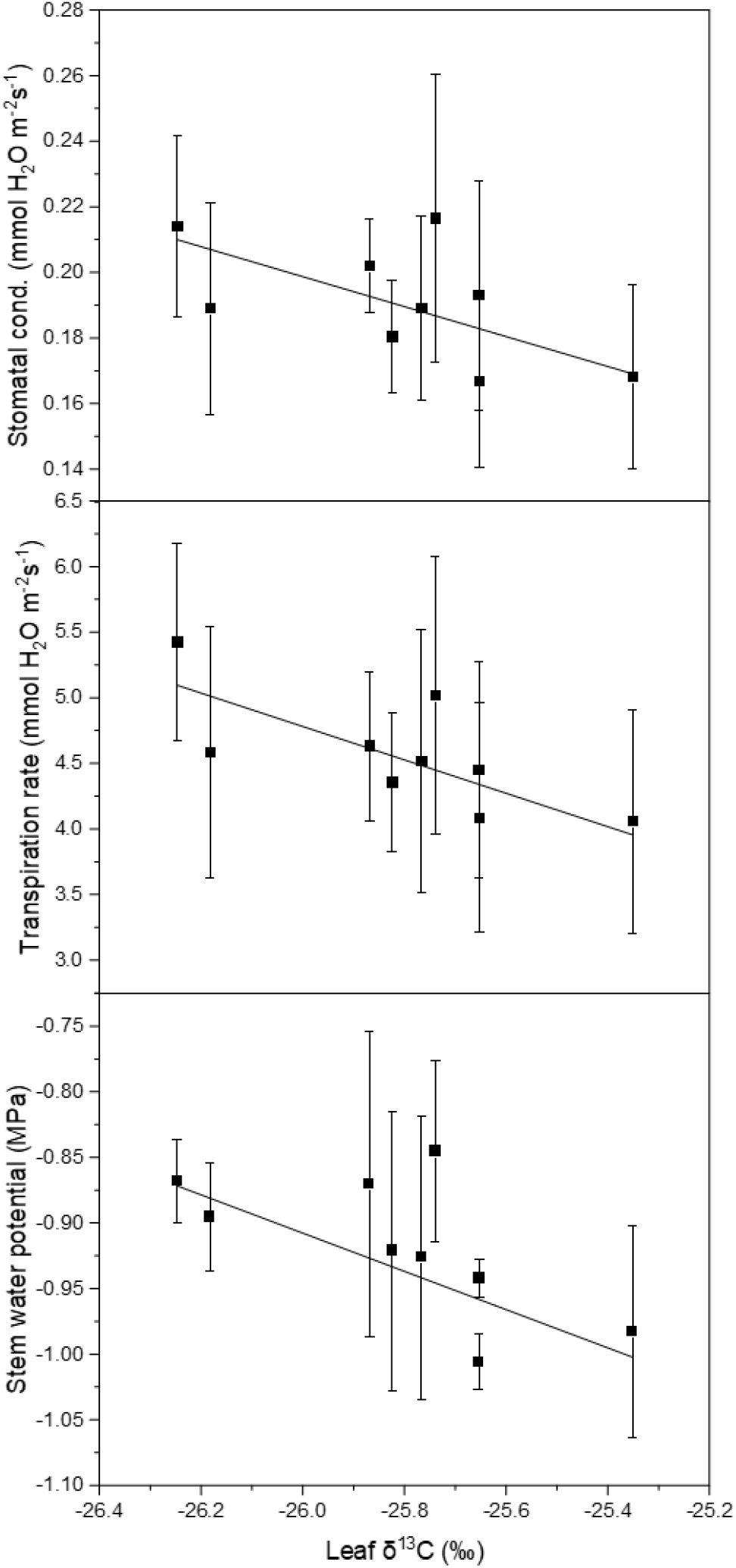
Stomatal conductance, transpiration rate, and stem water potential versus leaf δ^13^C (N=9) in 2018. Bars denote standard deviation of mean values for each replication.

## CONCLUSION

Rootstock genotype can affect photosynthesis rate, stomatal conductance and transpiration rate thereby influencing carbon and oxygen isotope composition of the scion. These responses are induced by changes to stem water potential which also affects overall tree vigor, a key trait imparted by rootstocks onto a scion. In this study, ‘Honeycrisp’ trees grafted on M.9 had elevated water limitations compared to more vigorous ‘Geneva’ rootstocks like G.87 and G.814. This is likely a result of elevated hydraulic resistance affecting tree growth and development. This condition will need more research in order to obtain a better understanding of the contributions how rootstocks affect water relations in the scion of apple and how it relates to tree vigor, productivity, and fruit quality.

## ACKNOWLEDGEMENTS

E.C.B and L.K. were supported by the USDA National Institute of Food and Agriculture Specialty Crop Research Initiative project “AppleRoot2Fruit: Accelerating the development, evaluation and adoption of new apple rootstocks” (2016-51181-25406). LK was also supported by the USDA National Institute of Food and Agriculture, Hatch/State project 1014919 and NC-140 Multistate Project “Improving Economic and Environmental Sustainability in Tree-Fruit Production Through Changes in Rootstock Use”, accession no. 1013833.

